# Structural basis for active-site probes targeting *Staphylococcus aureus* serine hydrolase virulence factors

**DOI:** 10.1101/2020.04.21.054437

**Authors:** Matthias Fellner, Christian S. Lentz, Sam A. Jamieson, Jodi L. Brewster, Linhai Chen, Matthew Bogyo, Peter D. Mace

## Abstract

*Staphylococcus aureus* is a major cause of infection in the community and in hospitals. Serine hydrolases play key roles in bacterial homeostasis, in particular biofilms. Activity-based profiling has previously identified a family of serine hydrolases, designated fluorophosphonate-binding hydrolases (Fphs), which contribute to virulence of *S. aureus* in the biofilm niche. Here we report structures of the putative tributyrin esterase FphF, alone and covalently bound by a substrate analog, and small molecule inhibitors that occupy the hydrophobic substrate-binding pocket. We show that FphF has promiscuous esterase activity. Building from this, we extended our analysis to the wider Fph protein family using homology modeling and docking tools. We predict that other Fph enzymes, including FphB which was linked directly to virulence, may be more specific than FphF. This study provides insight into Fph function and a template for designing new imaging agents, diagnostic probes, and inhibitors to treat *S. aureus* infections.

## 1. Introduction

*Staphylococcus aureus* populates mucosal tissues or skin on about 30% of the world’s population, and is a common cause of a variety of diseases ranging from local skin or soft tissue infections to invasive infections such as bacteremia, pneumonia or endocarditis (Tong et al., 2015a; Wertheim et al., 2005). Of particular concern are drug resistant forms such as Methicillin-resistant *S. aureus* (MRSA), which are a major cause of mortality (Turnidge et al., 2009). The increased occurrence of community-acquired antibiotic-resistant *S. aureus* strains is an especially major health threat, requiring urgent development of new diagnostic and therapy options (Tong et al., 2015b). Of particular clinical concern are also biofilm-based infections. Biofilms are surface-associated bacterial communities embedded in an extracellular matrix that provides a barrier to the host immune system and difficult to eradicate. Cells in a biofilm have been reported to be in a state of reduced metabolic activity and growth that causes tolerance to antibiotic treatment and can lead to chronic infections in the clinic (Lebeaux et al., 2014; Waters et al., 2016). In order to successfully target bacteria in this restrictive growth state, an increased understanding of the underlying bacterial physiology and biochemistry along with a thorough validation of any druggable enzyme in a biofilm environment is imperative.

Serine hydrolyses are one of the largest enzyme families in nature, they are highly druggable and may execute a plethora of different biological functions. Yet, the role of serine hydrolyses in bacterial homeostasis, survival at the host-pathogen interface or in biofilm-associated growth is not well explored. In order to mine this enzyme family for new targets in anti-virulence and anti-infectivity strategies, we have recently performed a cell-based chemical proteomics study in *S. aureus*, employing activity-based probes (ABPs) (Lentz et al., 2018). ABPs are functionalized enzyme inhibitors that can be used for selective labelling of active enzymes to characterize their physiological roles *in vitro* and *in vivo*. Target-specific probes can provide leads for small molecule inhibitors, enable direct analysis of drug efficacy and specificity and be used to create selective imaging agents. In our recent work, we used fluorophosphonate-probes to identify 12 serine hydrolyses targets that are enzymatically active under biofilm forming conditions. These enzymes include lipase1 and 2 (SAL1, SAL2) as well as 10 uncharacterized hydrolases that we termed fluorophosphonate-binding hydrolases (Fph) A-J (Lentz et al., 2018). A detailed investigation of one particular hydrolase, FphB, revealed that it is a new virulence factor facilitating infection in specific tissue sites in a mouse model of systemic infection (Lentz et al., 2018). As part of the initial studies, we identified selective inhibitors and fluorescent activity-bases probes for several Fph proteins (Chen et al., 2019; Lentz et al., 2018). Structural knowledge for these new potential virulence factors has the potential to establish a structure-function relationship for Fph proteins as well as create a template for further development of inhibitors and active site probes with increase potency and selectivity (Chen et al., 2019; Lentz et al., 2018). Based on their selective inhibition profile, biological relevance and accessibility to chemical probes, serine hydrolases appear promising target candidates. However, serine hydrolases are prevalent in both bacteria and host organisms. Therefore, understanding the structure and function roles of specific hydrolases in bacterial physiology or pathogenesis and how substrate selectivity is achieved, is crucial to enable strategies to target these enzymes.

In this study, we present the crystal structure and substrate specificity profile of one of the most abundant biofilm expressed Fph proteins, the putative tributyrin esterase FphF. This enzyme is a 29 kDa hydrolase encoded by a gene previously annotated as *estA*. In the UniProt database, this protein (Q2FUY3) is putatively annotated as Tributyrin esterase and has a GO-annotation as an S-formylglutathione hydrolase based on similarity with the human orthologue. Our biochemical and structural data suggest that FphF is a promiscuous enzyme, with broader substrate specificity for a wider range of hydrophobic acyl groups than other *estA* encoded serine hydrolases. In addition, our experimentally determined structure allowed us to model the nine remaining Fph proteins of unknown function. Furthermore, we obtained crystal structures of FphF in complex with a substrate and triazole urea-based inhibitors KT129 and KT130, enabling docking analysis of substrates and inhibitors for various Fph family members. These models provide a basis for understanding the active site environments of these enzymes that share a common fold but lack sequence identity. While our findings suggest that FphF is promiscuous, we predict that the majority of the other Fph proteins are more specific. Most Fph proteins show a tighter substrate binding site compared to FphF that include several residues with functional groups, suggestive of distinct substrate binding. Given the roles of other Fph proteins in virulence, they are likely ideal targets for imaging agents, diagnostic probes, inhibitors, and ultimately therapeutic agents.

## 2. Materials and methods

### FphF cloning, expression and purification for activity measurements

The full length *fph*F (currently annotated as *estA*, gene loci SAOUHSC_02962, NWMN_2528, SAUSA300_2564) was amplified from the *S. aureus* ATCC35556 genome using primers ATGA<u>GGATCCGCTT</u>ATATTTCATTAAACTATCA and <u>GAAACTCGAGTTAATCATTCACCATCCATGTT</u> that introduced XhoI and BamHI restriction sites, respectively. The PCR product was gel-purified and extracted before double-digestion with XhoI and BamHI-HF and dephosphorylation with Antarctic Phosphatase. The resulting gene fragment was ligated into XhoI- and BamHI-HF-double digested pET28a using T4 DNA ligase (NEB). The ligation mixture was transformed into NEB 5-α-competent *Escherichia coli*.

For expression of recombinant full length-FphF the pET28a-SAOUHSC02962 plasmid was transformed into BL21 (DE3) competent *E. coli* and grown on LB-Kanamycin (35 μg/mL) selection media. An overnight culture of the transformed bacteria in selection medium was diluted 1:250 into 500 mL LB-Kanamycin in a 2 L flask and grown at 37°C, 220rpm until the culture reached an OD_600_ of 0.5. Recombinant protein expression was induced by addition of 10 μM Isopropyl β- d-1-thiogalactopyranoside. Cultures were continued to shake at 27°C for 4 h, before cells were harvested by centrifugation (8000 g, 10 min, 4°C). The cell pellet was re-suspended in a small volume of LB medium, transferred to two 50 mL polypropylene tubes and centrifuged again (4000g, 10 min, 4°C). The supernatant was discarded and cell pellets were frozen at −80°C. Each pellet was re-suspended in 8 mL Lysis Buffer (50 mM NaH2PO4, 300 mM NaCl, 10 mM imidazole) and lysed by sonication. The lysate was cleared by centrifugation at 4,350 g, 30 min, 4°C and the supernatant was added to 1 mL Ni-NTA-Agarose. The sample was incubated and mixed by rotation at 4°C for 1-2 h. The resin was washed 3x with wash buffer (Lysis buffer with 20 mM imidazole) before His6-tagged protein was eluted with elution buffer (Lysis buffer with 250 mM imidazole) in 8 fractions of 1 mL. Eluates were pooled and concentrated 10-fold in 10,000 MWCO spin columns. 10 mL of 50 mM Tris-HCl, pH8.0, 300 mM NaCl, 20% glycerol were added and the samples were concentrated 10-fold. Samples were combined and protein concentration determined by OD280 measurements using the sequence-specific calculated extinction coefficient (E1% = 14.24).

### ABP-labeling of recombinant FphF

Recombinant FphF was diluted into PBS/0.01% SDS (50 nM) and was pre-incubated with JCP678 or DMSO at 37°C for 30 min before FP-TMR was added (1 μM final concentration) for fluorescent labelling of active protein at 37°C for an additional 30 min. After addition of 4x SDS-PAGE, the samples were boiled at 95 C for 10 min, cooled down and 35 ng of protein were analyzed by SDS-PAGE. The gel was scanned for TMR fluorescence on a Typhoon 9410 variable mode imager (λ_ex_= 380 nm, 580 BP filter). Subsequently, the gel was subjected to silver staining and photographed over a transilluminator.

### FphF enzymatic activity assays

The hydrolytic activity of purified recombinant FphF protein was tested using a series 4-Methylumbelliferyl(4-MU)-based fluorogenic substrates as described previously (Lentz et al., 2018). In brief, 0.3 μL of fluorogenic substrates (10 mM in DMSO) were added to the wells of an opaque flat-bottom 384 well plate. 30 μL of a 10 nM solution of FphF in PBS/0.01%TritonX-100 was added and fluorescence (λ_ex_ = 365 nm and λ_em_ = 455 nm) was read at 37°C in 1 min intervals on a Cytation 3 imaging reader (BioTek, Winooski, VT, USA) for 60 min. In the linear phase of the reaction (10 – 40 min) turnover rates were calculated using Gen5 software (BioTek) as RFU/min and were normalized by subtracting background hydrolysis rates measured for each substrate in reaction buffer in the absence of protein.

### FphF cloning, expression, and purification for crystallization

FphF DNA construct was designed based on UniProt sequence Q2FUY3 omitting the starting methionine with overhangs for ligation-independent cloning (Luna-Vargas et al., 2011). The construct was synthesized by Integrated DNA Technology (IDT) and cloned into modified pET28a-LIC vectors incorporating an N-terminal 6× His tag and a 3C protease cleavage site.

*E. coli* BL21(DE3) cells in 1 L cultures (1X Luria-Bertani with 50 μg/mL kanamycin) at 37 °C and 200 rpm shaking were grown until OD_600_ reached of 0.6. Cultures were induced with 0.2 mM Isopropyl β-d-1-thiogalactopyranoside and grown over night at 18 °C and 200 rpm shaking. Bacterial pellets were harvested via centrifugation, suspended in 50 mM Tris pH 8.0, 300 mM NaCl and stored at −20 °C.

For purification thawed suspended pellets were incubated for 30 min on ice with ~20 μg/mL lysozyme and ~4 μg/mL DNase. Cells were lysed via sonication (Sonifier Heat Systems Ultrasonics). FphF protein was initially purified by Ni^2+^ affinity chromatography (HIS-Select resin, Sigma-Aldrich) using an elution buffer containing 50 mM Tris pH 8.0, 300 mM NaCl, 300 mM Imidazole, 10% (v/v) glycerol and 10% (w/v) sucrose. Elution fractions were incubated with 3C protease and 2 mM DTT over night at 4 °C. FphF was further purified by size-exclusion chromatography using anion exchange (RESOURCE Q) and/or Superdex 75 or 200 Increase columns (GE Life Sciences). Anion exchange separated two FphF species of unknown difference, potentially representing two different dimer forms. Size-exclusion only showed a single peak. Both anion exchange peaks looked identical on SDS-PAGE analysis and both yielded crystals. The first eluted major peak (~80% of protein) was used for most experiments and resulted in the presented datasets. Either chromatography step or a combination appeared to result in crystallizable protein, the final column buffer used for crystal drops is given in the next section. Purified protein was either used directly for crystal drops or snap frozen in liquid nitrogen.

### FphF crystallization

FphF was broad screened for crystallization resulting in multiple hits with details given in supplemental table S1. After optimization, the following three datasets were obtained, fully refined and deposited.

For the FphF apo-form 0.2 μL ~8.5 mg/mL FphF (20 mM HEPES pH 7.5, 10 mM NaCl) were mixed with 0.1 μL FphF crystal seeds (in 54.4% Tacsimate pH 7.0, 0.1 M Bis-Tris propane pH 6.5, 8% Polypropylene glycol P 400) and 0.2 μL of reservoir solution. Sitting drop reservoir contained 50 μL of 2.8 M sodium acetate. Crystals were soaked for ~20 seconds in 75% reservoir solution and 25% ethylene glycol prior to freezing in liquid nitrogen.

For FphF KT129 bound 0.2 μL ~7.5 mg/mL FphF+KT129 (0.12 mM KT129, 12% DMSO, 18 mM HEPES pH 7.5, 8 mM NaCl) were mixed with 0.2 μL of reservoir solution. Sitting drop reservoir contained 50 μL of 0.2 M sodium citrate, 0.1 M Bis-tris propane pH 6.5, 20% w/v PEG 3350. Crystals were soaked for ~20 seconds in 75% reservoir solution and 25% glycerol prior to freezing in liquid nitrogen.

For FphF KT130 bound 0.25 μL ~6.6 mg/mL FphF+KT130 (0.2 mM KT130, 20% DMSO, 8 mM HEPES pH 7.5, 40 mM NaCl) were mixed with 0.2 μL of reservoir solution. Sitting drop reservoir contained 50 μL of 0.8 M sodium formate, 10% w/v PEG 8000, 10% w/v PEG 1000 and 0.1 M HEPES pH 7.0. Crystals were soaked for ~20 seconds in 75% reservoir solution and 25% glycerol prior to freezing in liquid nitrogen.

For FphF heptyl acyl bound 0.4 μL ~8.0 mg/ml FphF (10 mM HEPES pH 7.5, 10 mM NaCl) were mixed with 0.07 μL ligand solution (~0.5 mM 4-MU heptanoate in 100% DMSO) and 0.4 μL of reservoir solution. Sitting drop reservoir contained 50 μL of 0.8 M Sodium formate, 10 % w/v PEG 8000, 10 % w/v PEG 1000 and 0.1 M Tris pH 7.5. Crystals were soaked for ~20 seconds in 75% reservoir solution and 25% glycerol prior to freezing in liquid nitrogen.

### FphF data collection and processing

X-ray diffraction data were collected at the Australian synchrotron MX1 (Cowieson et al., 2015) and MX2 (Aragao et al., 2018) beamlines. Datasets were processed with xds (Kabsch, 2010), merging and scaling were performed using aimless (Winn et al., 2011). Phases were solved with Phenix Phaser molecular replacement (McCoy et al., 2007) using a model created with BALBES (Long et al., 2008). Model building and refinement were conducted in COOT (Emsley et al., 2010) and Phenix (Afonine et al., 2012). Ligand creation and restraint generation utilized Jligand (Lebedev et al., 2012) and eLBOW (Moriarty et al., 2009).

All FphF apo-protein crystal contained a three atom modification to Ser121. The electron density peak for the central atom ~2.4 Å from the serine OG atom always was significantly bigger than a water molecule would account for. No anomalous dispersion peak was observed in electron density maps at this site, ruling out a heavy metal. A covalent modification to Ser121 was rules out by mass spectrometry. Purified FphF and FphF crystals were subjected to mass spectrometry analysis, but no covalent post-translational modification could be detected. Sodium chloride was the only compound present in across all crystallization conditions, it was therefore modelled as a sodium ion with two water molecules. Two other α/β hydrolases with similar active site serine bound sodium ions from *Coxiella burnetii* (PDB ID 3TRD (Franklin et al., 2015)) and *Pseudomonas fluorescens* (3T4U unpublished) were identified using DALI (Holm, 2019) and rcsb.org (Berman et al., 2000) searches.

Modeling and refinement of the ligand bound KT129 and KT130 structures was complicated as multiple conformations appeared to exist in the same datasets. It appeared plausible that the apo-protein sodium form might be present at a low occupancy as well as two conformations of the ligands. In the KT129 bound structure the evidence for these other forms were not strong enough and only one KT129 conformation was modelled at an occupancy of ~74%. However, in the KT130 structure, a second conformation appeared to be present at a higher occupancy, especially in chain C where we modelled this conformation resulting in KT130 ligand occupancies of 55% A and 45% B. The heptyl acyl group also had hints of a possible second conformation in some chains but refinement of a two conformation of the ligand were not satisfying.

Statistics for the datasets are listed in Table S2 and S3. Structure figures, analysis and alignments were created with ChemSketch (ACD/ChemSketch, 2018), UCSF Chimera (Pettersen et al., 2004) and Procheck (Laskowski et al., 1993).

### Structure prediction and docking

Structure prediction was done by the I-Tasser (Yang et al., 2015) webserver using chain A of the FphF apo structure (PDB ID 6VHD). The only other input was sequences obtained from UniProt: FphA (UniProt ID Q2FVG3, predicted molecular weight 52.0 kDa); FphB (Q2FV90, 36.8); FphC (Q2FYZ3, 35.3); FphD (Q2G2D6, 33.2); FphE (Q2FV39, 31.0); FphF (Q2FUY3, 29.1); FphG (Q2G2V6, 28.4); FphH (Q2G025, 28.1); FphI (Q2G0V7, 27.4); FphJ (Q2FVA9, 21.8). The resulting models were manually inspected and compared to FphF. The best model was picked by examining the prediction of the core secondary structure elements, the location of the active site triad, as well as statistics provided by I-Tasser. Ligand creation, conversion and manipulation used the tools PRODRG (Schuttelkopf and van Aalten, 2004), Open Babel (O’Boyle et al., 2011) and COOT (Emsley et al., 2010). Molecular docking experiments were performed with GOLD (Jones et al., 1997). Default parameters were used if not stated otherwise with hydrogens added to the protein structure using the gold_serine_protease_VS template. The binding site was centered at the active site serine residue. For uncleaved ligands the active site serine was mutated to a glycine. For docking triazole inhibitors ligand flexibility options were utilized. For covalent docking the GOLD instructions were followed, adding a “link oxygen atom” to the ligands and defining the link between the active site serine and the link oxygen atom.

## 3. Results

### Overall structure of FphF

FphF crystallized in several different conditions (Table S1) in three distinct crystal forms. The overall oligomeric structure was nearly identical across the three crystal forms showing a tetramer (Figure 1A and 1B) either within the asymmetric unit or across symmetry mates. In solution, FphF appears to be a dimer of ~58 kDa based on gel-filtration experiments (Figure S1A). There was no significant difference between the FphF monomers (Cα RMSD of 0.25-0.35 Å between the chains 20) and the biological function of the oligomer remains uncertain, potentially contributing to overall stability. FphF is a member of the α/β hydrolase superfamily (Ollis et al., 1992), indicated by core 8 β strands connected by several α helices (Figure S2).

**Figure 1:**
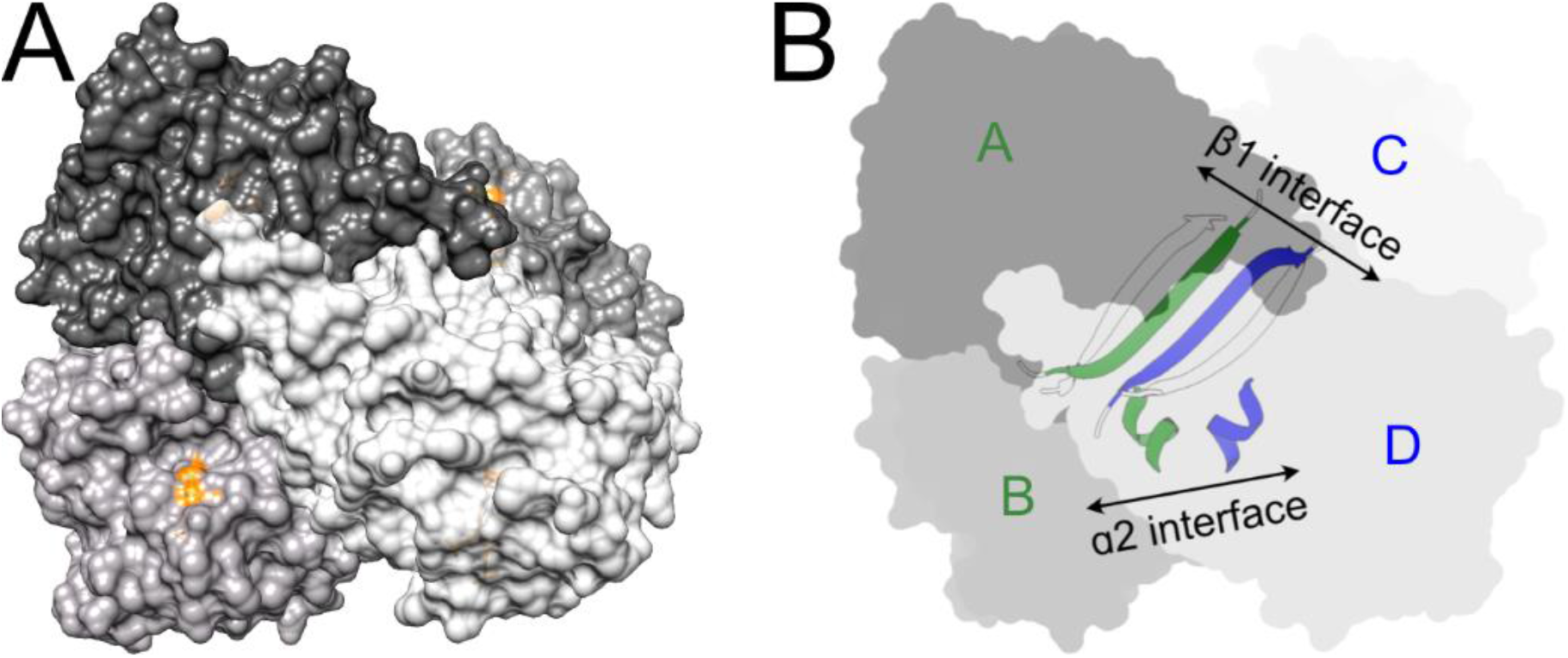
Overall structure of FphF. A) FphF tetramer (PDB ID 6VH9). Surface of four Chains are colored in different shades of grey with active site triad in orange. B) Two different dimer interfaces within the tetramer indicated by blue and dark green coloring of β1-β1 and α2-α2.

In comparing FphF to known structures, DALI (Holm, 2019) searches revealed 260 representative structures hits, but no structure with more than 30% sequence identity. The top DALI candidate (27% sequence identity) was tributyrin esterase (estA) from *Streptococcus pneumonia* (PDB ID 2UZ0 (Kim et al., 2008); sequence alignment Figure S3). EstA is a virulence factor, which induces inflammation by inducing NO production and cytokine secretion in macrophages (Kang et al., 2009). EstA has been shown to be active against tributyrin and various acetylated substrates (Kahya et al., 2017). EstA adopts the same dimer of dimer tetramer conformation as FphF and is also described as a dimer in solution (Kim et al., 2008). There are two significant dimer interfaces in both tetramers. One predominantly formed by an antiparallel association of the first β-strand β1, while the second one is formed around interactions of the second helix α2 (Figure 1B and S1B). The overall structural similarities suggest that *S. aureus* FphF and *S. pneumonia* EstA are related, however our following structure-function characterization show clear differences between the two enzymes.

### Active site of FphF

The FphF serine hydrolase catalytic triad consists of Ser121, His234 and Asp205 (Figure 2A and 2B). Ser121 is located on the connecting loop between β5 and α6 at the conserved GXSXG motif (Schrag and Cygler, 1997), which all Fph proteins contain. His234 sits on a loop between β8 and the conserved C-terminal helix α11 and Asp205 on a loop between β7 and α10. Across all FphF apo-protein crystal forms and all crystallization conditions, Ser121 is found proximal to a nearby three atom density, which we modelled as a sodium ion chelated by two waters. For details on this density assignment see methods section and Figure S4. We speculate that the sodium binding illustrates the reactivity of the Ser residue and is otherwise only a minor off pathway interaction and possibly a crystal artifact.

**Figure 2:**
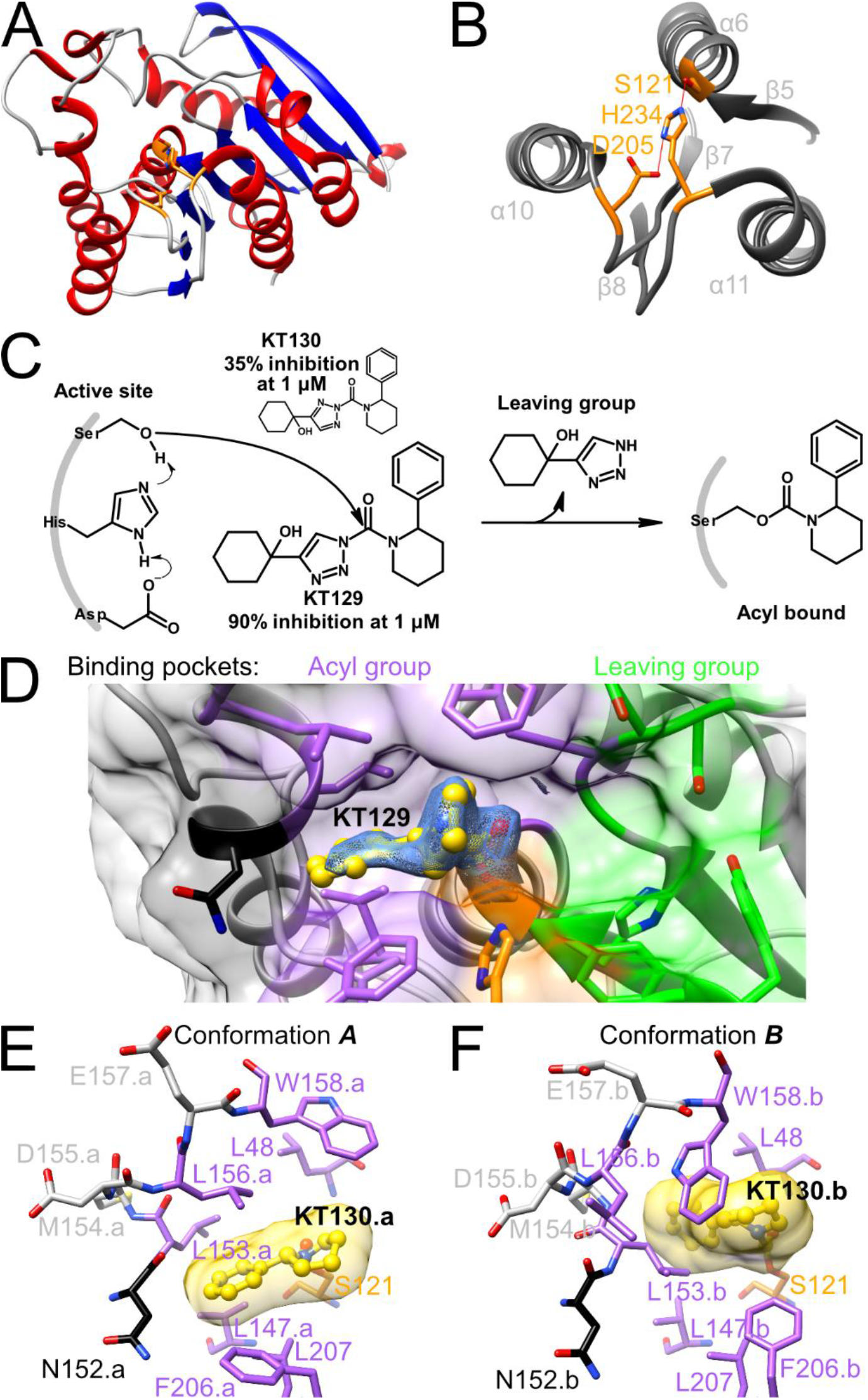
Covalent inhibitor bound FphF crystal structures. A) Ribbon representation of FphF monomer showing the location of the active site triad. β-strands in blue, α-helices in red, active site Ser-His-Asp triad in orange. B) Closeup of the active site triad. C) KT129 and KT130 mode of FphF inhibition, with % of inhibition at 1 μM (Chen et al., 2019). D) FphF KT129 bound structure in conformation ***A*** (6VHD) is shown with ligand carbon atoms in yellow, the active site triad in orange, the acyl binding pocket in purple, the Asn125 gate in black, the leaving group binding pocket in green with other residues in grey (this coloring scheme is used for all other figures). The *2FO−FC* map for KT129 is shown as blue mesh at 1 σ. D) KT130 bound model (6VHE) in conformation ***A*** with transparent ligand surface. E) KT130 bound model in conformation ***B*** with transparent ligand surface.

### Covalent inhibitor bound structure of FphF

We previously identified several 1,2,3-triazole urea based inhibitors which preferentially targeted FphF (Chen et al., 2019). In particular, two isomers, KT129 and KT130, which differed in the constitution of the triazole urea linkage had potencies that differed by an order of magnitude (Figure 2C). We determined co-crystallized structures of FphF with KT129 and KT130. As expected based on the enzymatic mechanism (Figure 2C), the triazole leaving group that distinguishes KT129 and KT130 was absent, leaving the identical 2-phenylpiperidine-1-carbonyl moiety covalently attached to the terminal side chain oxygen atom of Ser121 revealing the acyl binding pocket of FphF (Figure 2D, E and F).

In both structures, all protomers of the tetramer contained electron density (Figure 2D) consistent with the ligand within a hydrophobic pocket alongside the active site. Two distinct conformations of the acyl group of KT129 and KT130 are observed. The predominate conformation “***A***” (Figure 2D and 2E) was clearly present in all protomers. Several hydrophobic residues surrounded the phenylpiperidine Leu48, Val147, Leu153, Leu156, Trp158, Phe206 and Leu207. Asn152 gates the entrance to this pocket, however its sidechain is turned towards the surface with its uncharged backbone interacting with the phenylpiperidine. Compared to binding pocket in the apo-protein, there are only small differences in which Trp158 and Phe206 side chains are slightly displaced from the active site upon ligand binding, suggesting a predominantly direct, non-induced fit of the ligand. In only some protomer active sites of the tetramer, there is electron density evidence of a possible second lower occupied “***B***” conformation (Figure 2F) which induces more changes at the active site. ***A*** appears to represent the major and biological relevant binding form, as the backbone and side chains of the apo-protein structure already resemble this conformation.

One outstanding question that can be partially addressed by the structures is the binding-site of the triazole leaving group of KT129 and KT130. This part of the ligand will bind at the surface exposed area adjacent to Ser121 (Figure 2C; green residues). In all structures, 2 or 3 water molecules form a hydrogen bonding network within a small binding pocket that includes Ser49, Ser50 and His120. This pocket represents the key binding site for polar elements within the leaving group. Nearby sidechains of Asp235, Tyr236, Trp239, adjacent to the active site H234 on the C-terminal helix α11, could also play important roles.

### FphF substrate profile and substrate bound structure

To establish how the active site observations from the inhibitor bound structures relate to enzymatic function, with respect to other Fph proteins, as well as EstA, we determined the substrate specificity profile of FphF. We also were able to determine the crystal structure of the protein bound to a preferred model substrate.

In order to confirm whether FphF has esterase activity, we used a fluorescent fluorophosphonate-probe (FP-TMR) that covalently binds to the active site serine of alpha-beta hydrolases. We previously used this probe to label FphF along with the other newly identified Fph enzymes in live *S. aureus* (Lentz et al., 2018). The probe labelled a single protein of the expected molecular weight of the recombinant protein including the N-terminal His6 tag (Figure 3A). In addition, we previously identified the sulfonyl fluoride JCP678 as a selective inhibitor of FphF in *S. aureus*. Pre-treatment of the purified protein with this covalent inhibitor, completely blocked labelling by the FP-TMR probe (Figure 3A). These results confirm that the labelling properties of the recombinant FphF protein are the same as native protein under physiological conditions in the cell. Next, we tested the substrate preference of FphF using a panel of commercially available fluorogenic substrates (Figure 3B). We found that the protein cleaved esterase substrates, but was unable to process phosphate, phosphonate or glycosidic substrates. FphF showed a promiscuous specificity profile as it is able to cleave hydrophobic lipid substrates with acyl chain lengths ranging from C2 to C10 with the highest activity for C7 (Figure 3B). To better understand the reason for this broad selectivity, we determined the crystal structure of FphF in complex with the preferred C7 model substrate 4-Methylumbelliferyl (4-MU) heptanoate (Figure 3C). In our structure, we were able to capture the heptyl acyl moiety of the substrate covalently linked to Ser121 in all FphF chains. Interestingly, the active sites were orientated very similar to conformation ***A*** of the inhibitor structures. The side chain orientations in the acyl binding pocket also matched the inhibitor bound state, with the exception of Phe206, which is disordered and can adopt several orientations between different protomers of the tetramer, with some facing away from the ligand (Figure 3C). Although at a lower resolution than the inhibitor bound structures, the substrate bound structure confirms that the hydrophobic binding pocket identified in the inhibitor structures is used to bind ester substrates.

**Figure 3:**
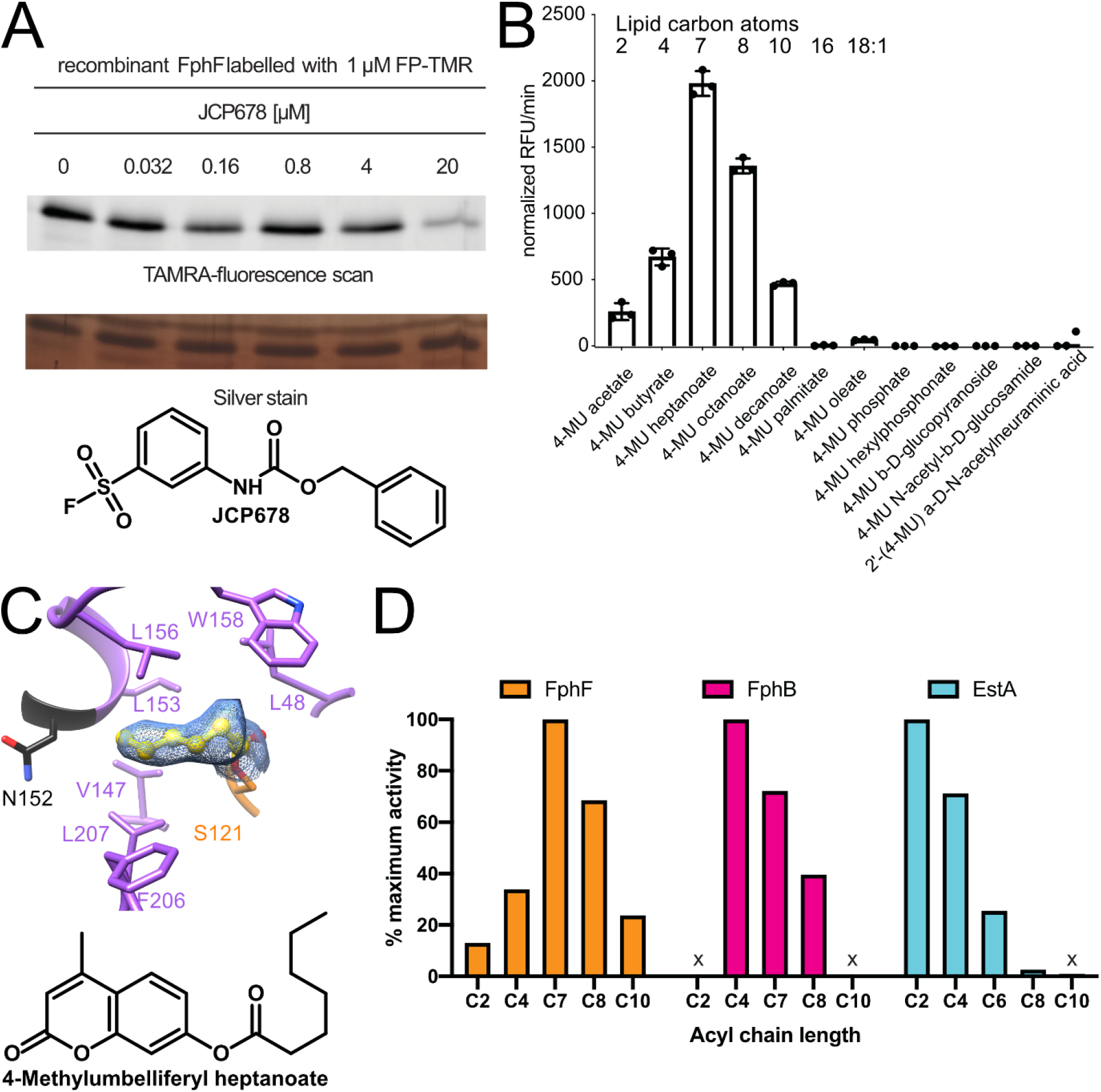
FphF enzymatic activity profile and substrate bound structure. A) Purified recombinant FphF was pretreated with different concentrations of the selective inhibitor JCP678 before labelling with the fluorescent ABP FP-TMR. B) Assessment of the substrate specificity profile using a library of 4-MU based fluorogenic substrates. The graph shows the turnover rates for each substrate as relative fluorescence units (RFU)/min and depicts the means +/− standard deviation of 3 independent reactions. The experiment was repeated twice with similar results. C) Covalent substrate (4-MU heptanoate) bound FphF crystal structure (PDB ID 6WCX). The *2FO−FC* map for the heptyl acyl intermediate is shown as blue mesh at 1 σ. D) the relative enzymatic activity of FphF, FphB and EstA as assessed in a series of fluorogenic carboxylic acid ester substrates of different acyl chain length (C2 – C10). The activity data of FphF and those reported previously for FphB (Lentz et al., 2018) and EstA (Kahya et al., 2017) were normalized by setting activity data measured for the preferred substrate to 100% and the data of the other substrates as the percentage of maximum activity. X denotes inactivity of the enzyme against the indicated substrate.

These findings prompted us to compare FphF with FphB, the only other Fph protein for which we have previously established a substrate profile (Lentz et al., 2018) (Figure 3D). FphB and FphF share only 21% amino acid sequence identity (sequence alignment Figure S3) and are predicted to have the same overall protein fold. The demonstrated carboxylic acid esterase activity of FphF is similar to FphB (Lentz et al., 2018). Like FphF, FphB efficiently processes hydrophobic lipid substrates but has a narrower specificity such that it is unable to cleave C2 and C10 lipids and has a preference for C4 substrates (Lentz et al., 2018). EstA on the other hand, had a preference for an even shorter chain length with a C2 acetate preference (Kahya et al., 2017), showing decreasing activity for C4, C6 and C8, and no activity for C10 (Figure 3D). We then proceeded using modeling and docking tools to understand the structural basis for differences in the selectivity profiles,

### Comparison of FphF, FphB and EstA using docking studies

We used FphF as a template to model FphB using I-Tasser (Yang et al., 2015), which combines the FphF input structure with up to nine additional structures from the protein data bank (Berman et al., 2007). We then performed docking studies into the FphF, EstA crystal structures and the FphB predicted model.

First, we docked ligands into the FphF-KT129 crystal structure. Covalent docking the acyl group of KT129 into the protein structure after removing the ligand gave a similar structural model for the ligand as observed in the crystal structure (Figure 4A). The hydrophobic pocket is identified by the docking program, with the ligand adopting slightly different conformations. Docking the full length KT129 inhibitor (Figure 4B) showed a similar fit for the acyl group, with the remaining part of the ligand occupying the leaving group binding pocket. The triazole nitrogen atoms are surface exposed, with the carbon atoms of the triazole ring facing the protein. The lone hydroxyl group is effectively stabilized by hydrogen bonds with Ser50, His120 and Tyr236. Docking of KT130 resulted in unique orientations for the hydroxyl group induced by the triazole ring (Figure 4B). One nitrogen of the triazole ring must face the protein, resulting in a tilt of the remaining ligand in this binding pocket. This results in the hydroxyl group forming hydrogen bonds with Ser49 and its adjacent backbone. 1,4-isomers of inhibitors such as KT129 were more potent against FphF in general when compared to the 2,4-isomers found in KT130 (Chen et al., 2019), suggesting that a nitrogen facing the protein is unfavorable for binding in the leaving group pocket. Comparison of the FphF-substrate structure (Figure 4C) with the docked full length 4-MU heptanoate shows the acyl group again occupying this pocket. For the full length ligand the acyl is also docked nearly perfectly on top of residue 121. The acyl group binding pocket is fully occupied by the C7 chain with several side chains clearly defining the space. However, the different orientation of the chain between the structure and docked ligand, as well as the access from the surface demonstrates that a longer chain length could also bind, explaining the substrate profile of FphF. The leaving group binding pocket is also identified for the 4MU moiety. The leaving group binding pocket is widely surface exposed and could accompany various functional groups and sizes of substrates. Together, these data support a promiscuous substrate profile of FphF.

**Figure 4:**
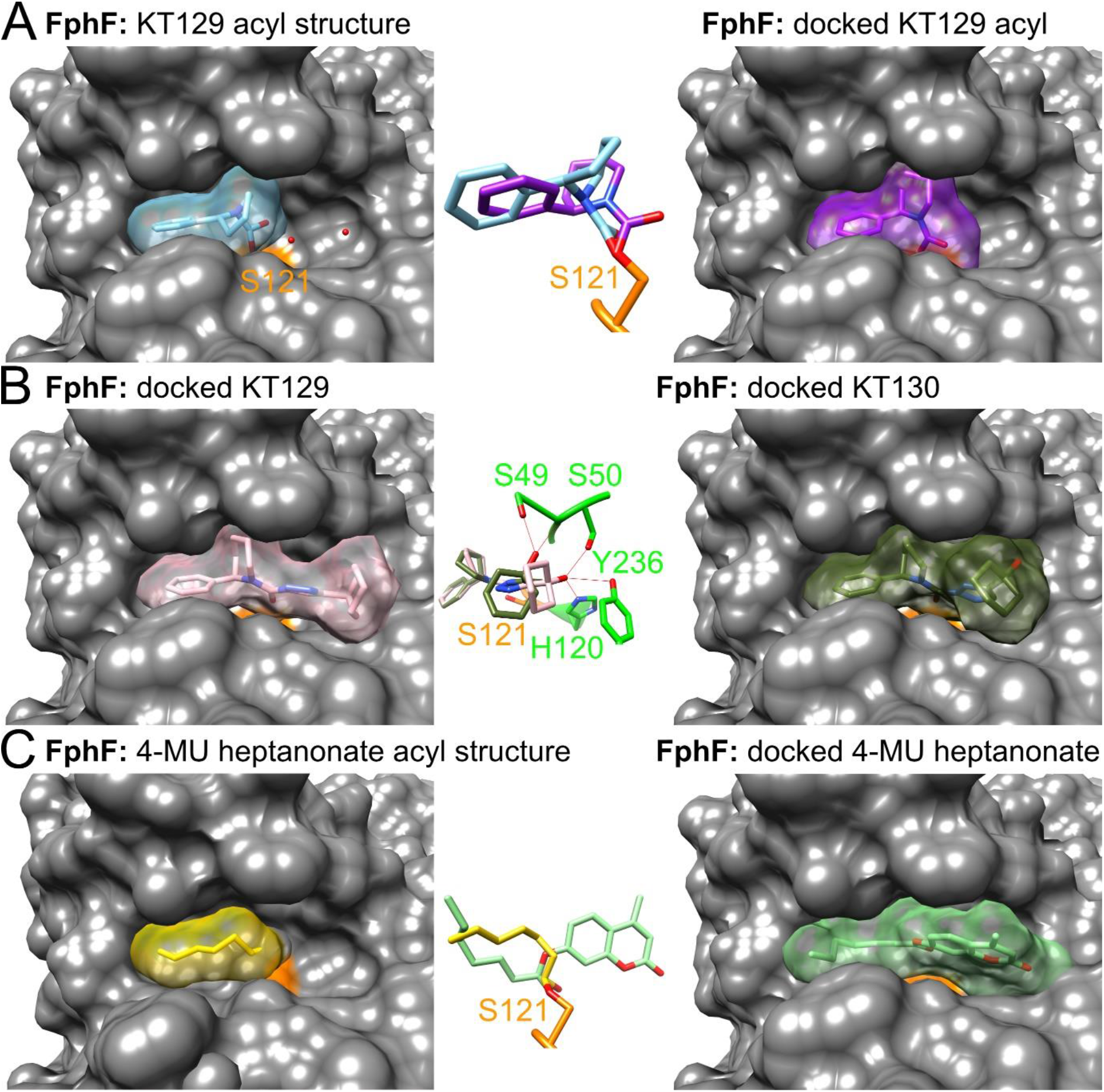
FphF docking studies A) KT129 acyl structure and docking comparison. FphF-KT129 structure (PDB ID 6VHD); protein surface in grey, except for active site Ser in orange; ligand and ligand surface in light blue. Water molecules in the leaving group pocket illustrated as red spheres. FphF; covalent acyl group of KT129 in purple docked into 6VHD. Middle insert shows overlay of the two. B) Full length KT129 and KT130 docking comparison. KT129 ligand and ligand surface in pink, KT130 ligand and ligand surface in dark green docked into 6VHD-S121G. Middle insert shows overlay of the two with differing hydrogen bonding residues of the ligand hydroxyl group in green. C) Heptyl acyl structure and docking comparison. FphF-substrate bound structure (6WCX); heptyl acyl ligand and ligand surface in yellow. FphF; full length 4-MU heptanoate in green docked into 6VHD-S121G. Middle insert shows overlay of the two.

We next compared the modelled structure of FphB to the structure of FphF. Fph proteins all belong to the α/β hydrolase superfamily (Ollis et al., 1992) and all of the predicted Fph protein models show this fold containing the core β strands and several of the conserved helices, including FphB. Comparison of the FphF structure active site triad and the corresponding residues in the FphB model demonstrate similar positions when aligning the full-length protein. This indicates a correctly folded active site in FphB, giving confidence to the prediction (Figure 5A). In the predicted model of FphB, the acyl binding pocket is not surface exposed. A loop containing Met225 (FphB numbering), in combination with a loop containing Lys43 between the FphB unique N-terminal helices, folds onto the active site (Figure 5B). The FphB Met225 loop is an extension of the 152-158 active site FphF loop. Not only is the FphB loop longer, the equivalent residues to 152-158 of FphF differ in FphB constricting the binding pocket even further. Although the FphB acyl binding pocket is buried, a clear pocket is defined by several hydrophobic residues (Figure 5C). The docking suggests that the covalent bound butanoyl of the preferred substrate 4-MU butyrate fits into this pocket. However, the acyl group of KT129 would likely not fit, confirming observations that KT129 and inhibitors with a similar sized acyl group are very poor inhibitors of FphB (Chen et al., 2019). Docking of the butanoyl moiety (Figure 5C and 5D) to the preferred model substrate of FphB shows that there is not much additional space for a longer lipid chain. This is in line with the substrate preference for a C4 chain length (Lentz et al., 2018) (Figure 3D). Docking the acyl group of AA395, a potent triazole inhibitor of FphB (Chen et al., 2019) (Figure 5E and 5F) also fits well into the binding pocket, filling up its entire space. Furthermore, for FphB, the leaving group binding pocket is blocked by several residues and an opening of this pocket by the Lys43 and Met225 loops is likely required to provide access to the active site serine. These findings further support an active site in FphB that has a narrower specificity compared to FphF.

**Figure 5:**
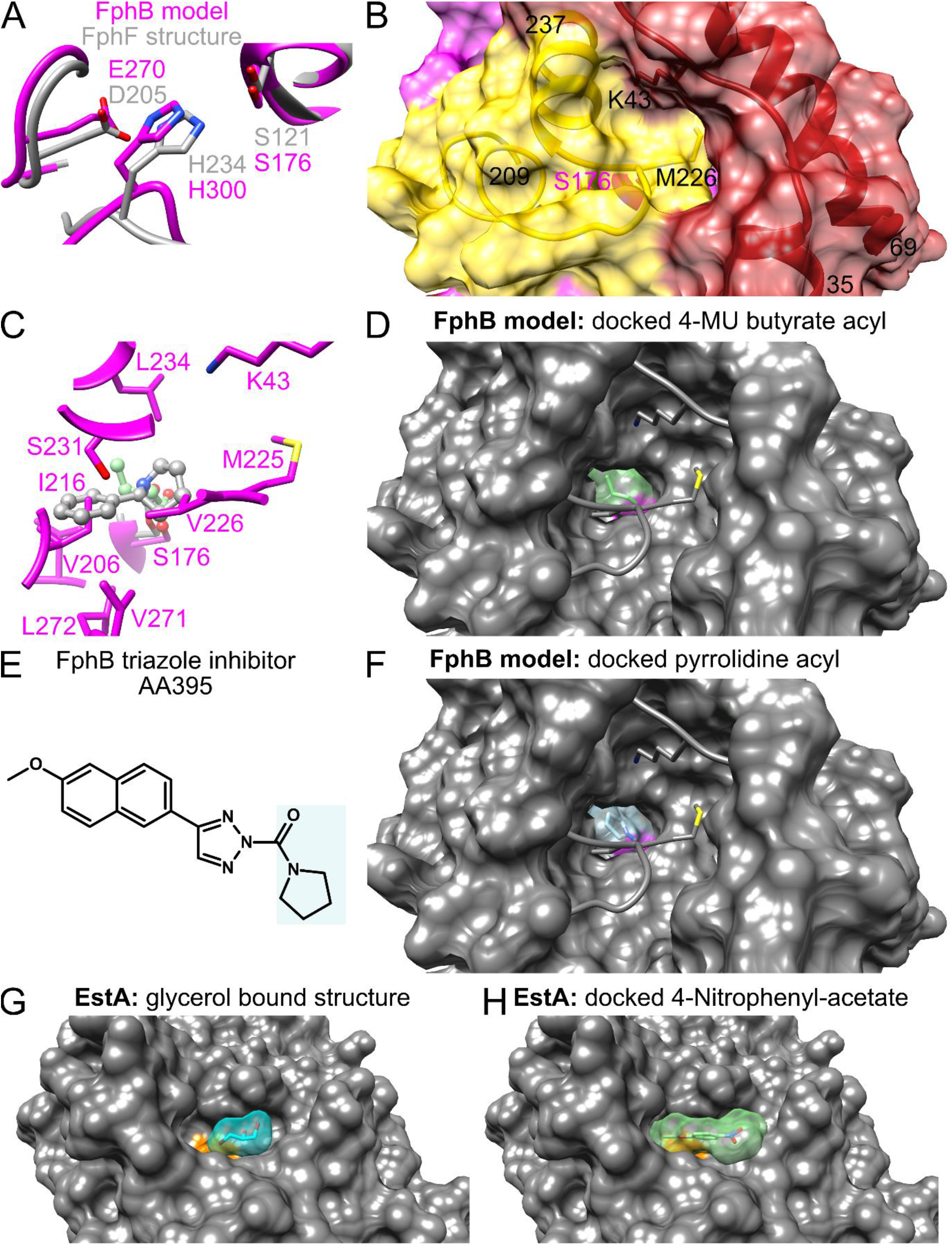
FphB and EstA docking studies. A) FphB predicted active site triad in magenta compared to FphF in grey based on a Cα alignment. B) FphB predicted protein surface 1-69 in dark red with side chain of K43 shown; 209-235 in yellow with side chain of M225 shown; remaining residues in magenta with buried active site S176 shown. C) Acyl binding pocket of FphB in magenta with ligands from FphF-KT129 acyl bound and docked covalent bound butanoyl of preferred substrate 4-MU butyrate. D) FphB surface representation of butanoyl bound. Ligand and ligand surface in green with S176 in magenta. Surfaces of loop around K43 and M226 are not shown. E) Chemical structure of the potent FphB inhibitor AA395. F) Same depiction as D with covalent docked acyl group of AA395. G) EstA structure (PDB ID 2UZ0); protein surface in grey, except for active site Ser in orange. Leaving group bound glycerol molecule in cyan. H) EstA; full length preferred substrate 4-Nitrophenyl-acetate in green docked into 2UZ0-S121G.

To gain additional insight into the substrate and leaving group binding pockets of the Fph hydrolases we compared the reported structure of EstA, which has a glycerol bound at this position (Kim et al., 2008) (Figure 5G). Docking the preferred model substrate 4-Nitrophenyl-acetate into EstA (Figure 5H) confirms that the leaving group sits at the position of the glycerol. The acetate acyl also fits nicely into the acyl binding pocket of EstA, which is narrow and formed by several hydrophobic residues around the active site Ser. Taken together, we can identify clear distinctions in the structures of FphF, FphB and EstA. While all act on hydrophobic acyl groups, their substrate specificities appear to differ significantly with FphF being the most promiscuous and FphB being the most specific. EstA likely has intermediate specificity between the two, but mainly functions as a deacetylase (Kahya et al., 2017).

### Comparison of FphF and FphB with other Fph proteins

Similar to FphB we used the FphF crystal structure, in combination with available data bank entries, to predict the structure of all 10 Fph proteins (Figure 6). While our models predict that the general protein fold is conserved across all the Fph proteins, alignment in Clustal Omega (Sievers et al., 2011) indicated that Fph proteins have overall low sequence similarities, with most having under 20% identity. We found secondary structure diversity in areas between the core β strands, at several loops and α-helices. Overall, the C-terminus is structurally conserved and always contains a terminal α-helix following the active site histidine loop. The N-terminus shows some variability before the first core β strand. Compared to FphF, some of the Fphs may have additional (FphA) or fewer (FphC) core β strands, which could influence their oligomerization. FphB and FphD stand out with predicted additional N-terminal extensions that contain two to three α-helices specific to these proteins. These helices may fold back onto the active site and close the binding pocket, also potentially introducing additional residues for distinct chemistry. However, this conformation is only seen for FphB and not FphD in the predicted models.

**Figure 6:**
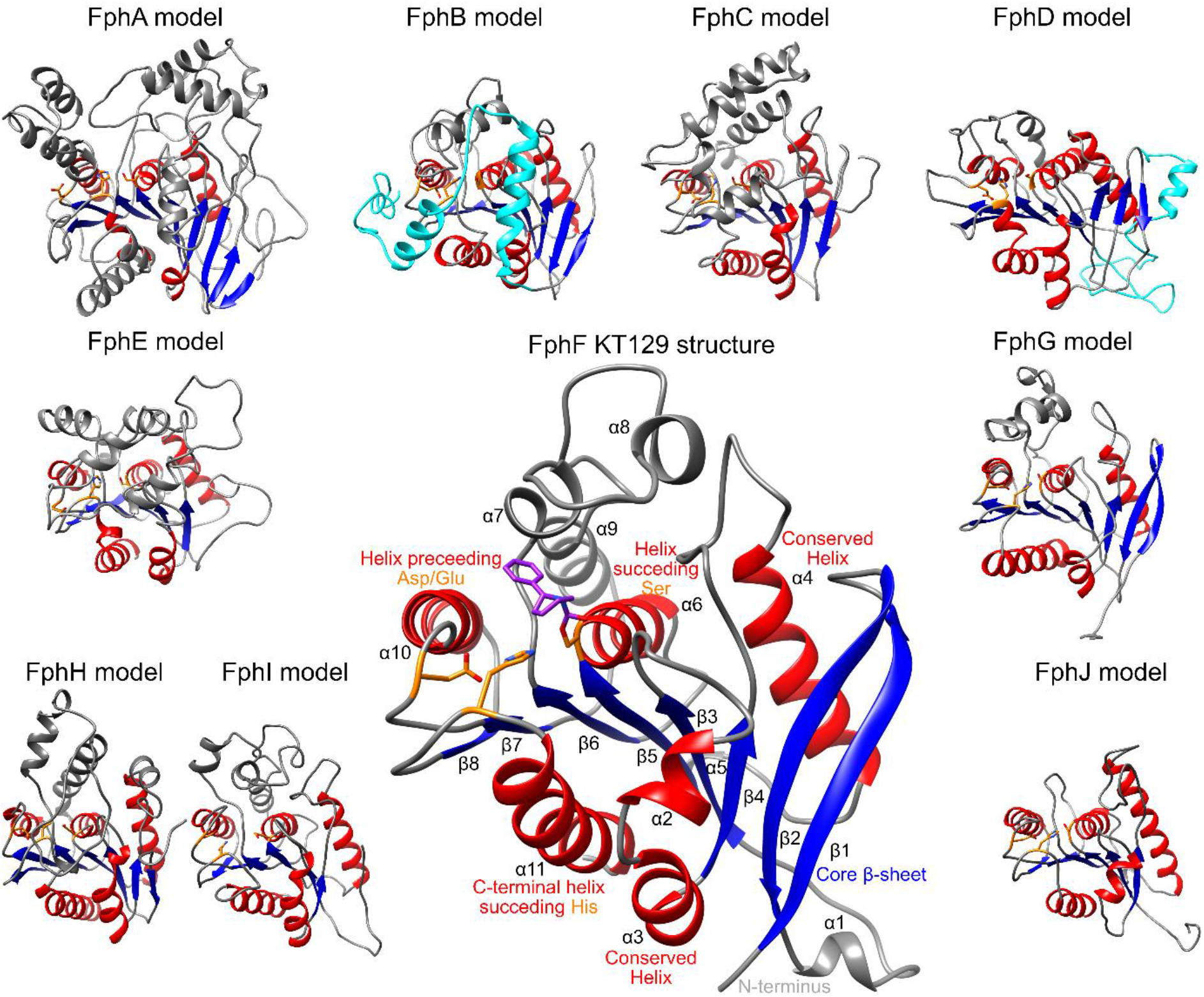
Predicted structures of Fph proteins. Structure of FphF KT129 bound in comparison to predicted models of the other Fph proteins. Figures based on full length Cα alignment. Core β-sheet in blue, conserved α-helices in red, active site Ser-His-Asp/Glu triad in orange. N-terminal extension in FphB and FphD in cyan. Secondary structure labelling in the FphF structure, bound KT129 in purple.

When comparing the predicted active sites of all the Fph proteins, it appears that in contrast to FphB and FphF, most Fph proteins may not prefer hydrophobic acyl groups. The acyl binding pocket of FphA, FphC, FphD, FphE, FphG and FphI have at least one and often multiple charged side chains pointing towards the acyl binding pocket. The remaining FphH and FphJ do not show many residues in the pocket, instead having a very open acyl binding pocket. FphA (Figure 7) introduces hydrophobic and charge residues in a narrow but surface exposed pocket. This suggests that FphA may have a highly selective substrate binding mechanism. This could possible explain why our triazole inhibitor screening only identified modest inhibitors for FphA. FphG (Figure 7) has the greatest diversity of residues predicted to be at the acyl pocket. Overall, FphB and FphF appear to have the most similar hydrophobic acyl binding pockets with all the other Fph proteins having a large variety of binding pockets. The leaving group binding pocket also seem to be as diverse across the family. For example, FphG has a diverse range of charged residues in this region, while FphJ has a clearly hydrophobic pocket (Figure 7). The overarching conclusions from these modeling studies is that there is little evidence for redundancy with respect to the Fph proteins. The overall low sequence identity is reflected in their predicted active sites, suggesting a wide range of substrate specificities, and thus potential functions.

**Figure 7:**
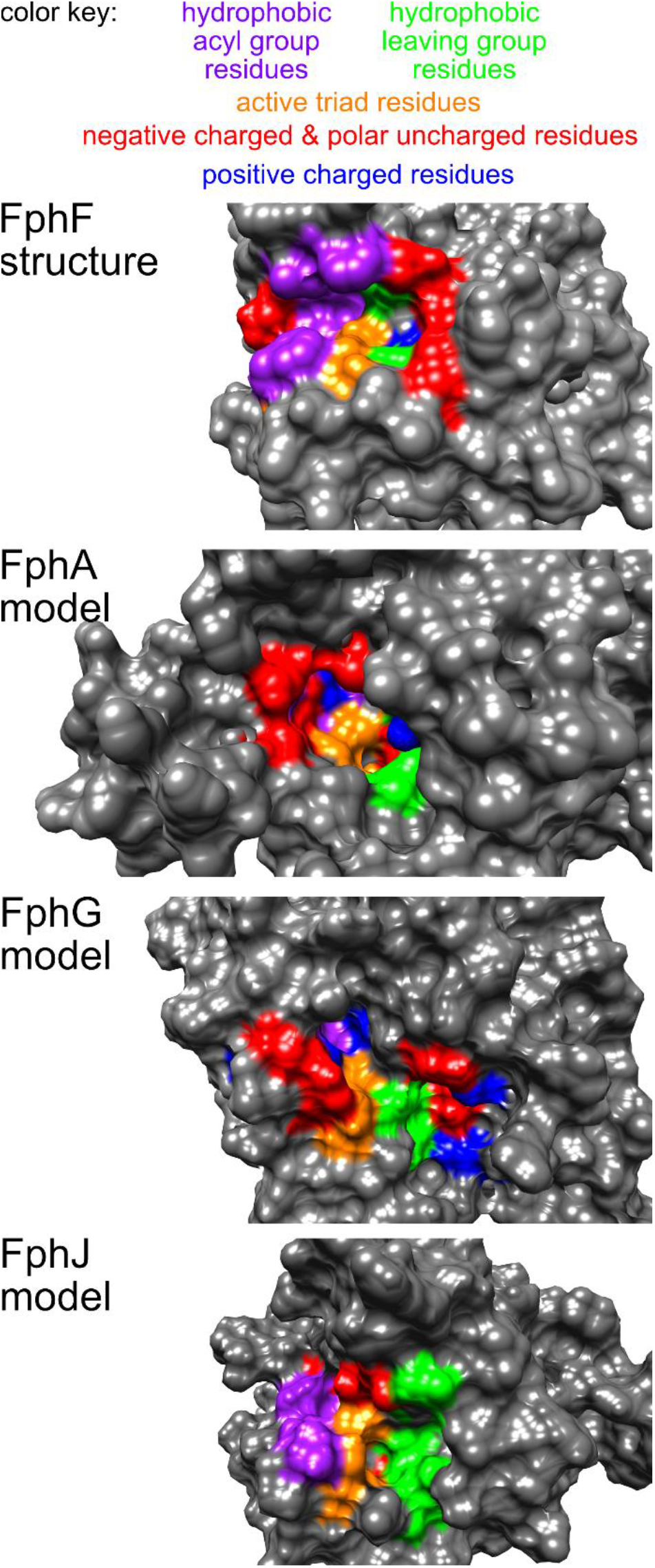
Fph proteins active sites. FphF active site compared to examples of predicted active sites of Fph proteins.

## 4. Discussion

Given that serine hydrolases can be effectively targeted by chemical probes, they are a promising targets for diagnosis, therapeutics and monitoring of treatment of *S. aureus* infections. One major problem that remains is the specific targeting of a particular enzyme over related enzymes in the host and the pathogen itself. A clear understanding of the structure-function relationships of these highly related enzymes is necessary to overcome this problem. Our structural characterization of FphF presented here demonstrates how we can gain insight into the specificities of serine hydrolases of *S. aureus*. While FphF, the related virulence factor FphB and the closest homolog from *S. pneumoniae* virulence factor, EstA (PDB ID 2UZ0 (Kim et al., 2008)) all appear to be related, there are several key differences. The acyl binding pockets of all three proteins are hydrophobic, confirming the observed substrate preference for extended lipid acyl chains. The different chain length preference is clearly related to the size of the acyl binding pocket in the structure of these three proteins. FphF is promiscuous, with a bigger acyl binding pocket, and is able to accommodate a wide range of hydrophobic substrates chain lengths. EstA acyl binding pocket is substantially smaller, with several hydrophobic residues defining a pocket optimal for C2 binding. Not surprisingly, EstA was shown to deacetylate O-acetylated sialic acid, allowing the bacteria to process acetylated mucins (Kahya et al., 2017). The FphB predicted model suggests an even higher level of specificity as additional loops can fold onto the acyl and leaving group binding pocket and potentially introduce additional residues for defining substrate preferences.

Although, our findings do not rule out that FphF is able to deacetylate native substrates, our data suggests that alternative, preferred physiological substrates of FphF may exist. Because the function of FphF is not likely to be similar to EstA, we propose to change the gene annotation of FphF and genes with high sequence identity to *fphF* from *estA*. While FphF can process a variety of lipid substrates, our structure analysis suggests that FphF may be able to process a variety of hydrophobic acyl groups of ester substrates, potentially including branched lipids and aromatic chains. Substrate specificity determinants may exist on both sides of the scissile bond and further studies are necessary to probe the leaving group binding pocket in greater detail.

When searching for possible homologs of FphF in *Homo sapiens* the only notable structure similarity and sequence identity hit (sequence alignment Figure S3) was human esterase D (ESD, also known as S-formylglutathione hydrolase, 3FCX (Wu et al., 2009)), but the overall identity was only at 23%. Optimization of targeting FphF or FphB could include parallel studies with ESD, as well as EstA to demonstrate specificity.

Even with knowledge of the substrate binding pocket of FphF, it is not entirely straightforward to predict the substrate-binding pockets of the other Fph proteins. In some cases, sequence alignment is poor in these regions and exact prediction of the active site pockets unreasonable. This is not unusual for a large superfamily such as the α/β hydrolases, where active sites are tuned to determine substrate specificity. Using the knowledge from our FphF structures, combined with available structure prediction and docking tools, we can begin to define similarities and differences between FphF and the other Fph proteins. This analysis suggests that the Fph enzymes have specific roles in *S. aureus* facilitated by their different predicted active site environments. Expanding the structure-function relationship, optimally using additional native substrates, will be crucial to understand the specific roles of each Fph protein. This will allow us to better map the repertoire of serine hydrolases required for *S. aureus* homeostasis and virulence.

### Significance

Around a third of healthy humans are carriers of *Staphylococcus aureus*, such that they have the bacteria on their skin without any active infection or disease. Despite being harmless in most individuals, *S aureus* can cause pathogenic infections. It often exists in biofilms in human tissue, resulting in a biomolecular matrix that is largely impermeable to the immune system and many traditional antibiotics. The increased occurrence of community-acquired antibiotic-resistant *S. aureus* strains, often linked to biofilm formation, is a major health threat, requiring urgent development of new diagnostic and therapy options. Serine hydrolases are a large family of enzymes that play key roles in bacterial homeostasis and survival at the host-pathogen interface during infection. They play a role in biofilms, contributing to the difficulty of achieving effective treatment. This makes serine hydrolases promising new anti-virulence and anti-infectivity targets. Activity-based profiling recently identified a family of serine hydrolases, designated fluorophosphonate-binding hydrolases (Fphs), which contribute to virulence of *S. aureus* in the biofilm niche. Here we report a structure-function characterization of one of these serine hydrolases, FphF, expressed during biofilm forming conditions. We determined that FphF is a promiscuous enzyme, able to cleave hydrophobic lipid substrates with a range of acyl chain lengths. Using this newly acquired structural data and similarities among the Fph family, we show that other Fph proteins, including FphB which was linked directly to virulence, may have a more well-defined substrate specificity. Our structural and biochemical studies confirm that FphF is distinct from previously characterized enzymes, making it an important reference enzyme in the serine hydrolase superfamily. Overall, our results provide insight into the specificity and the mechanism of substrate and chemical probe binding to Fph. This information will aid in future efforts to targeting serine hydrolase virulence factors from *S. aureus* and other related bacteria.

## Acknowledgments

This work was supported by a University of Otago Health Sciences Postdoctoral Fellowship [HSCDPD1703 to M.F.], a postdoctoral research fellowship of the German Research Foundation (DFG) to C.L., a China Scholarship Council fellowship No. 201604910274 (to L.C.), a Youth Innovation Promotion Association of the Chinese Academy of Sciences grant No.2017329 (to L.C.).

