## Supplemental information for "Structural basis for active-site probes targeting *Staphylococcus aureus* serine hydrolase virulence factors"

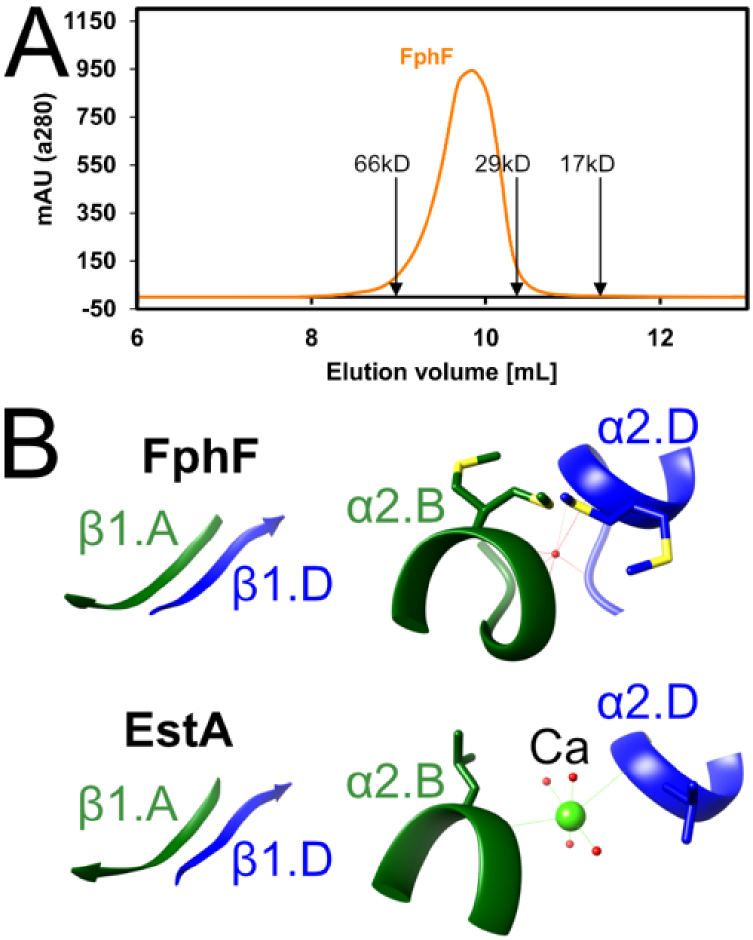


**Figure S1: Dimer of FphF.** A) A280 Superdex 75 chromatograph comparison of 58 kDa FphF dimer (29 kDa monomer) compared to molecular weight standard proteins bovine serum albumin 66 kDa; carbonic anhydrase 29 kDa and diubiquitin 17 kDa. C) Comparison of dimer interfaces between FphF and EstA (PDB ID 2UZ0). At the α2 interface FphF with alternate conformations of Met56 and chelated water compared to EstA with an interfacing Calcium ion.

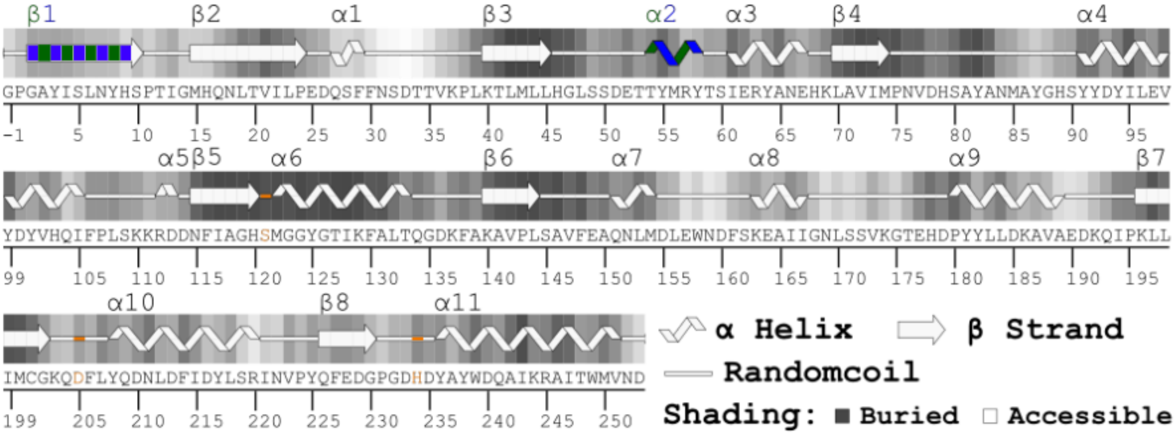


**Figure S2: Sequence and corresponding secondary structure of FphF.** Active site triad in orange. Two different dimer interfaces within the tetramer indicated by blue and dark green coloring of β1-β1 and α2-α2.

```
FphF S. aureus      MAYISLNY----------------------------------------------------  8
FphB S. aureus      --MRKKWSTLAFGFLVAAYAHIRIKEKRSVKSYMLEQGIRLSRAKRRFMYKEEAMKALEK  58
EstA S. pneumoniae  MAVMKIEY----------------------------------------------------  8
ESD  H. sapiens     MALKQISSNKCFGGLQKVFEH---------------------------------------  21
                        .

FphF S. aureus      -----------------HSPTIGMHQNLTVILPEDQSFFNSDTTVKPLKTLMLLHGLSSD  51
FphB S. aureus      MAPQTAGEYEGTNYQFKMPVKVDKHFGSTVYTVND------KQDKHQRVVLYAHGGAWFQ  112
EstA S. pneumoniae  -----------------YSQVLDMEWGVNVLYPDANRVEE--PECEDIPVLYLLHGMSGN  49
ESD  H. sapiens     -----------------DSVELNCKMKFAVYLP-------PKAETGKCPALYWLSGLTCT  57
                                         :. .    *                   .*    *

FphF S. aureus      ETTYMRYTSIERYANEHKLAVIMPNVDHS----------AYANM-------------AYG  88
FphB S. aureus      DPLKIHFEFIDELAETLNAKVIMPVYPKI------------------------PHQDYQAT  149
EstA S. pneumoniae  HNSWLKRTNVERLLRGTNLIVVMPNTSNG----------WYTDT-------------QYG  86
ESD  H. sapiens     EQNFISKSGYHQSASEHGLVVIAPDTSPRGCNIKGEDESWDFGTGAGFYVDATEDPWKTN  117
                    .   :    ..         *: *

FphF S. aureus      HSYYDYILEVYDYV-HQIFP-LSKKRDDNFIAGHSMGGYGTIKFALTQGDKF----AKAV  142
FphB S. aureus      YVLF---EKLYHDLL-----NQVADSKQIVVMGDSAGGQIALSFAQLLKEKHIVQPGHIV  201
EstA S. pneumoniae  FDYYTALAEELPQVLKRFFPNMTSKREKTFIAGLSMGGYGCFKLALTTN-RF----SHAA  141
ESD  H. sapiens     YRMYSYVTEELPQLINANFPV---DPQRMSIFGHSMGGHGALICALKNPGKY----KSVS  170
                    .  :    :    :          . .   : * * **   :  *     :.

FphF S. aureus      PLSAVFEAQNLMDLEWNDFSKEAIIGNLSSVK---GTEHD----PYYLLDKAVAEDKQIP  195
FphB S. aureus      LISPVLDATMQHPEIPDYLKKDPMVGVDGSVFLAEQWAGDTPLDNYKVSP-INGDLDGLG  260
EstA S. pneumoniae  SFSGALSFQNFSPESQNLGSPAYWRGVFGEIR---DWTTS----PYSL-ESLAKKSDKKT  193
ESD  H. sapiens     AFAPICN---PVLCPWG---KKAFSGYLGTDQ--SKWKA------YDATHLVKSYPGSQL  216
                     ::   .         .        *  .                *

FphF S. aureus      KLLIMCGKQD-FLYQDNLD---FIDYLSRINVPYQFEDGPGD---HDYAYWDQAI-KRAI  247
FphB S. aureus      RITLTVGTKE-VLYPDALN---LSQLLSAKGIEHDFI--PGYYQFHIYPVFPIPERRRFL  314
EstA S. pneumoniae  KLWAWCGEQD-FLYEANNL---AVKNLKKLGFDVTYSHSAGT---HEWYYWEKQL-EVFL  245
ESD  H. sapiens     DILIDQGKDDQFLLDGQLLPDNFIAACTEKKIPVVFRLQEGY--DHSYYFIATFI-TDHI  273
                     :    * .: .*              .   .   :    *    * :           :

FphF S. aureus      TWMVND--------  253
FphB S. aureus      YQVKNIIN------  322
EstA S. pneumoniae  TTLPIDFKLEERLT  259
ESD  H. sapiens     RHHAKYLNA-----  282
```

**Figure S3: Sequence alignment.** Alignment of FphF (UniProt ID Q2FUY3); FphB (Q2FV90); EstA (A0A0H2UNZ8) and ESD (P10768). Active site triad highlighted.

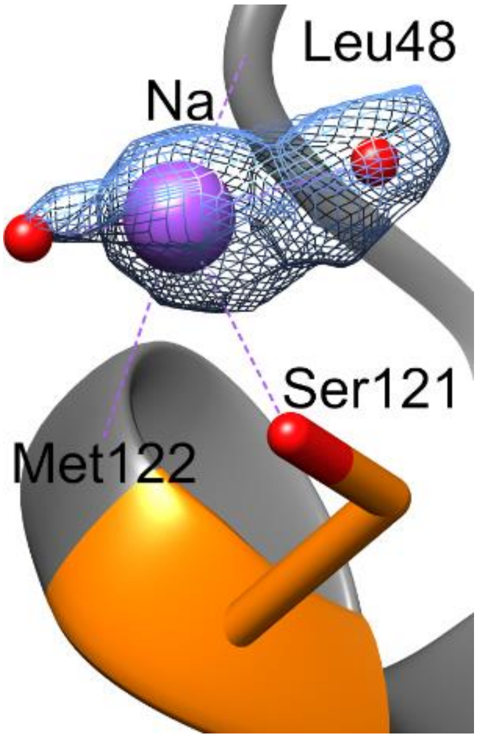


**Figure S4: Sodium binding in the FphF apo-protein**. The *2FO−FC* maps for sodium atom (purple sphere) and chelating water molecules (red sphere) are shown as blue meshes at 1 σ (PDB ID 6VH9). Sodium coordination to Ser121, backbone of Met122 and backbone of Leu48 illustrated as purple dashes.

**Table S1: Crystallization conditions of FphF.**

Highest diffracting crystal in Å. Crystal form 2 showed significant anisotropy and translational noncrystallographic symmetry.

| (Å) | Compound 1 | Compound 2 | Compound 3 |
|---|---|---|---|
| **Crystal form 1*** | | | |
| 1.7 | 2.8 M Sodium acetate | | |
| 1.9 | 0.2 M Trisodium citrate | 0.1 M Bis-tris propane pH 6.5 | 20% w/v PEG 3350 |
| 1.9 | 0.8 M Sodium formate | 0.1 M Tris pH 7.5 or HEPES pH 7.0 | 10% w/v PEG 8000 10% w/v PEG 1000 |
| 2.2 | 0.7-0.8 M Sodium formate | 0.1 M Tris pH 7.5 or HEPES pH 7.0 | 25% w/v PEG MME 2000 |
| 2.2 | 1.5 M $Li_2SO_4$ | 0.1 M Tris pH 8.0 or 8.5 | |
| ~4 | 2.0 M $(NH_4)_2SO_4$ | 0.1 M Sodium cacodylate pH 6.5 | 0.2 M NaCl |
| **Crystal form 2**** | | | |
| 2.1 | 40-75% Tacsimate | 0-0.1 M Bis-tris propane pH 6.5-8.0 | 0-8% w/v Polypropylene glycol |
| 2.6 | 0.8 M Sodium formate | 0-0.1 M Tris pH 7.5 | 25% w/v PEG MME 2000 |
| 2.7 | 0.8-1 M Sodium formate | 0-0.1 M Sodium cacodylate pH 6.8 or HEPES pH 7.0 | 7-10% w/v PEG 8000 7-10% w/v PEG 1000 |
| 2.9 | 0.2 M Trisodium citrate | 0.1 M Bis-tris propane pH 6.5 | 20% w/v PEG 3350 |
| 3.0 | 1.4 M Disodium malonate | 0.1 M Bis-tris propane pH 7.0 | |
| ~8 | 0.2 M Ammonium acetate | 0.1 M HEPES pH 7.5 | 25% w/v PEG 3350 |
| **Crystal form 3***** | | | |
| 3.1 | 2.7-2.8 M Sodium acetate | 0-0.1 M Sodium cacodylate pH 6.8 M or Tris pH 7.5 | |
| 3.2 | 0.8 M Sodium formate | 0-0.1 M Tris pH 7.5 or HEPES pH 7.0 | 10% w/v PEG 8000 10% w/v PEG 1000 |
| 3.3 | 0.7-0.9 M Sodium formate | 0.1 M Sodium cacodylate pH 6.4 or Tris pH 7.5 or 8.5 | 25% w/v PEG MME 2000 |
| 3.6 | 60% Tacsimate | 0-0.1 M Bis-tris propane pH 7.0 | 0-0.1 M $(NH_4)_2SO_4$ or Disodium malonate |
| 3.8 | 1.4-2.4 M Disodium malonate | 0-0.1 M Bis-tris propane pH 7.0 | |
| 3.8 | 1-1.4 M Trisodium citrate | 0.1 M Bis-tris propane pH 7.0 or Sodium cacodylate pH 6.5 or HEPES pH 7.0 | |
| ~4 | 0.2 M Trisodium citrate | 0.1 M Bis-tris propane pH 6.5 | 20% w/v PEG 3350 |
| ~5 | 0.8 M Sodium formate | 0.1 M Tris pH 7.5 | 25% w/v PEG MME 2000 |
| ~7 | 0.2 M Ammonium acetate | 0.1 M HEPES pH 7.5 | 25% w/v PEG 3350 |
| ~9 | 1.8 M $Li_2SO_4$ | 0.1 M Tris pH 8.8 | |

*Crystal form 1: $P6_1\,2\,2$; a, b, c (Å); α, β, γ (°) = ~87, 87, 454; 90, 90, 120; 4 chains

**Crystal form 2: $P6_2\,2\,2$; a, b, c (Å); α, β, γ (°) = ~88, 88, 222; 90, 90, 120; 2 chains

***Crystal form 3: $P6_4\,2\,2$; a, b, c (Å); α, β, γ (°) = ~88, 88, 110; 90, 90, 120; 1 chain

**Table S2: FphF data collection and processing**

Values for the outer shell are given in parentheses.

| | apo | KT129 bound | KT130 bound | heptyl acyl bound |
|---|---|---|---|---|
| Diffraction source | Australian synchrotron MX2 | Australian synchrotron MX2 | Australian synchrotron MX1 | Australian synchrotron MX1 |
| Wavelength (Å) | 0.954 | 0.954 | 0.954 | 0.954 |
| Detector | DECTRIS EIGER X 16M | DECTRIS EIGER X 16M | DECTRIS EIGER X 9M | DECTRIS EIGER X 9M |
| Space group | P $6_1$ 2 2 | P $6_1$ 2 2 | P $6_1$ 2 2 | P $6_1$ 2 2 |
| a, b, c (Å) | 87.1, 87.2, 453.6 | 87.2, 87.2, 455.2 | 87.1, 87.1, 454.9 | 87.0 87.0 454.7 |
| α, β, γ (°) | 90, 90, 120 | 90, 90, 120 | 90, 90, 120 | 90, 90, 120 |
| Resolution range (Å) | 49.16 – 1.71 (1.74 – 1.71) | 49.28 – 1.98 (2.02 – 1.98) | 49.24 – 1.94 (1.98 – 1.94) | 49.19 – 2.89 (3.07 – 2.89) |
| Total No. of reflections | 3,064,967 (144,842) | 1,980,193 (107,296) | 3,049,489 (145,969) | 217,394 (26,252) |
| No. of unique reflections | 112,281 (5,332) | 73,040 (4,307) | 77,616 (4,466) | 23,922 (3,641) |
| Completeness (%) | 99.9 (97.8) | 99.9 (98.5) | 100.0 (99.5) | 99.4 (97.4) |
| Redundancy | 27.3 (27.2) | 27.1 (24.9) | 39.3 (32.7) | 9.1 (7.2) |
| $\langle I/\sigma(I) \rangle$ | 16.5 (1.5) | 16.7 (1.5) | 23.1 (2.2) | 7.7 (1.6) |
| $CC_{1/2}$ | 0.999 (0.581) | 0.999 (0.711) | 1.000 (0.804) | 0.988 (0.524) |
| $R_{merge.}$ | 0.128 (3.155) | 0.151 (2.676) | 0.152 (1.957) | 0.223 (1.137) |
| $R_{p.i.m.}$ | 0.025 (0.605) | 0.029 (0.535) | 0.024 (0.342) | 0.069 (0.380) |

**Table S3: FphF structure solution and refinement**

Values for the outer shell are given in parentheses.

| | apo | KT129 bound | KT130 bound | heptyl acyl bound |
|---|---|---|---|---|
| Resolution range (Å) | 43.57 – 1.71 (1.73 – 1.71) | 49.28 – 1.98 (2.00 – 1.98) | 49.24 – 1.94 (1.96 – 1.94) | 49.19 – 2.89 (2.96 – 2.89) |
| Final $R_{cryst}$ | 0.177 (0.316) | 0.176 (0.301) | 0.168 (0.272) | 0.225 (0.305) |
| Final $R_{free}$ | 0.201 (0.342) | 0.213 (0.333) | 0.209 (0.332) | 0.261 (0.272) |
| Protein residues | 1020 | 1020 | 1020 | 1020 |
| Ligands | 4 (Na) | 4 (KT129) | 4 (KT130) | 4 (heptyl acyl) |
| Water | 457 | 258 | 499 | 17 |
| R.m.s. deviations | | | | |
| Bonds (Å) | 0.009 | 0.010 | 0.007 | 0.007 |
| Angles (°) | 1.190 | 1.399 | 0.948 | 0.947 |
| Average *B* factors (Å$^2$) | 39.5 | 46.9 | 38.0 | 43.9 |
| Ligands | 42.4 | 56.0 | 46.0 | 42.4 |
| Water | 45.0 | 48.3 | 40.9 | 36.0 |
| Ramachandran plot | | | | |
| Most favored (%) | 96.2 | 96.3 | 96.7 | 96.2 |
| Outlier (%) | 0 | 0 | 0 | 0 |
